# A Differential Equation for Mutation Rates in Environmental Coevolution

**DOI:** 10.1101/2021.03.19.436176

**Authors:** C E Neal-Sturgess

## Abstract

In their paper Natural selection for least action (Kaila and Annila 2008) they depict evolution as a process conforming to the Principle of Least Action (PLA). From this concept, together with the Coevolution model of Lewontin, an equation of motion for environmental coevolution is derived which shows that it is the time rate (frequency) of evolutionary change of the organism (mutations) that responds to changes in the environment. It is not possible to compare the theory with viral or bacterial mutation rates, as these are not measured on a time base. There is positive evidence from population level avian studies where the coefficient of additive evolvability (Cav) and its square (IA) change with environmental favourability in agreement with this model. Further analysis shows that the time rate of change of the coefficient of additive evolvability (Cav) and its square (IA) are linear with environmental favourability, which could help in defining the Lagrangian of the environmental effects.

## Background

Since the publication of “On the Origin of Species by Natural Selection” by Charles Darwin in 1859, there have been a number of attempts to link it to other scientific principles, notably the Principle of stationary Action, known popularly as “Least Action” [Nahin 2004]. In their paper Natural Selection for Least Action (Kaila and Annila 2008) depict evolution as a process conforming to the Principle of Least Action (PLA). This paper, although not giving any experimental evidence, shows that evolution, if conforming to the Second Law of Thermodynamics, will follow a trajectory of maximum Entropy production, which conforms to the PLA. To demonstrate this, they rewrote the Gibbs-Duhem relationship in terms of all possible states, to give a differential equation of evolution. This is a convincing argument as biology at root is a physical process, as expounded by Schrodinger 1944, and all physical processes are governed by the Second Law of thermodynamics.

It was difficult to reconcile evolution and the original Gaia hypothesis due to haemostasis [Lovelock 1979], however a later variant of Gaia called “Coevolution” [Lewontin 1983] allows reconciliation with Gaia. Lewontin’s description of Coevolution as two, coupled, differential equations to express the interaction between the organism (O) and the environment (E) as:

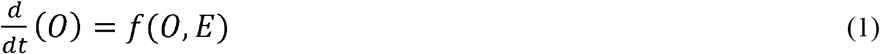

and

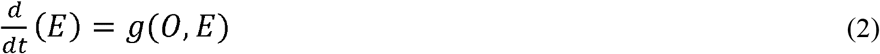

was an interesting development. It should be noted that in the literature coevolution can either be species coevolution and environmental evolution, this paper is solely concerned with environmental coevolution. Levin and Lewontin 1985 stated that solving these differential equations would be very difficult.

Taking the idea that the second law can be rewritten as an equation of motion [Kaila and Annila 2008] and if Lagrangian’s can be defined as:

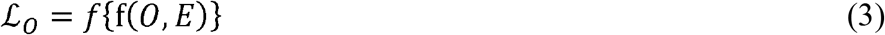

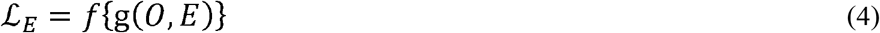

Then to find the unique trajectory in evolution-space time [Neal-Sturgess 2021] by using the calculus of variations through Euler-Lagrange [Goldstein 1969] to find an equation of motion for evolution gives:

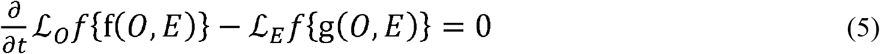

Hence:

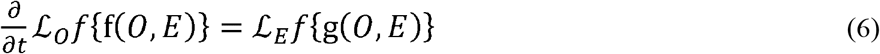

The Lagangians will be difficult to define, but it is possible that they will contain a potential term proportional to the environment, and a kinetic term proportional to the organisms muatation rate. Therefore this equation gives an partial differential equation for environmental coevolution and shows that the time rate of organism change,i.e. the rate of viral mutation or evolution rate, (the left hand side) is proportional to the magnitude of the change occurring in the environment (the right hand side) which is in agreement with some early evidence [Husby, Gross, Gianepp, Franks] dealt with below.

Much has been said and published about “Herd Immunity”, largely by politicians and journalists, however the scientific view is very different [Aschwanden 2020]. There is considerable difficulty calculating the proportion of the population with herd immunity necessary to suppress a virus, and in fact no natural or vaccine induced herd immunity has ever eradicated a virus; influenza is an example [Shao et.al 2017, Plans-Rubio 2012]. Plans-Rubio 2017 examined I_c_% which is the indicator of herd immunity from 1918 to 2010, where it is concluded that much higher levels of herd immunity are necessary to suppress viruses than have been achieved so far. There is a tension here between the present theory and the concept of herd immunity, for as the pressure on the virus occurs, either by natural or vaccine induced herd immunity, equation 6 still says that mutation rates will increase, leading to a scenario of constant catch-up.

To look for empirical evidence of equation 8 the field can be broadly classified as 1) viral mutations, 2) bacterial mutations and 3) evolutionary population level studies (considered by Levin and Lewontin as possibly the only level at which their equations may be solved). Considering viral mutation rates first, there is a considerable number of investigations reported in the literature [Sanjuan et.al. 2010. Regoes et.al 2012, Peck et.al. 2015, Sanjua & Domingo-Calap 2015, Martinez-Padilla et.al. 2017. Peck & Lauring 2018, Duffy 2018]. However, a problem exists when trying to compare the reported “viral mutation rates” with the analysis conducted here. Equation 6 is a “rate of change” with respect to time, whereas typically the viral mutation rates are reported as substitution’s/nucleotide/cell (s/n/c), which should not be confused with rates of evolutionary change [Peck & lauring 2018]. Although there is some evidence of a relationship between the two [Peck & Lauring 2018] see Fig 3, it is likely that any such relationship will probably be a function of virus type and the mutation mechanism. Pleck et.al. 2015 state that Neutral Theory [Kimura 1983] posits that the evolutionary rate increases linearly with the mutation rate, which is interesting in that equation 6 could, with appropriate constants. apply to both viral rates and evolutionary rate. Although there is clearly a relationship between evolutionary rate and mutation rate there does not appear to be any systematic investigations of viral mutation rates or evolutionary rate and the environment. However, there seems to be a general conclusion that viral mutation rates do respond to selective environmental pressures from the environment, which could be in agreement with the differential equation derived here.

**Fig. 2.**
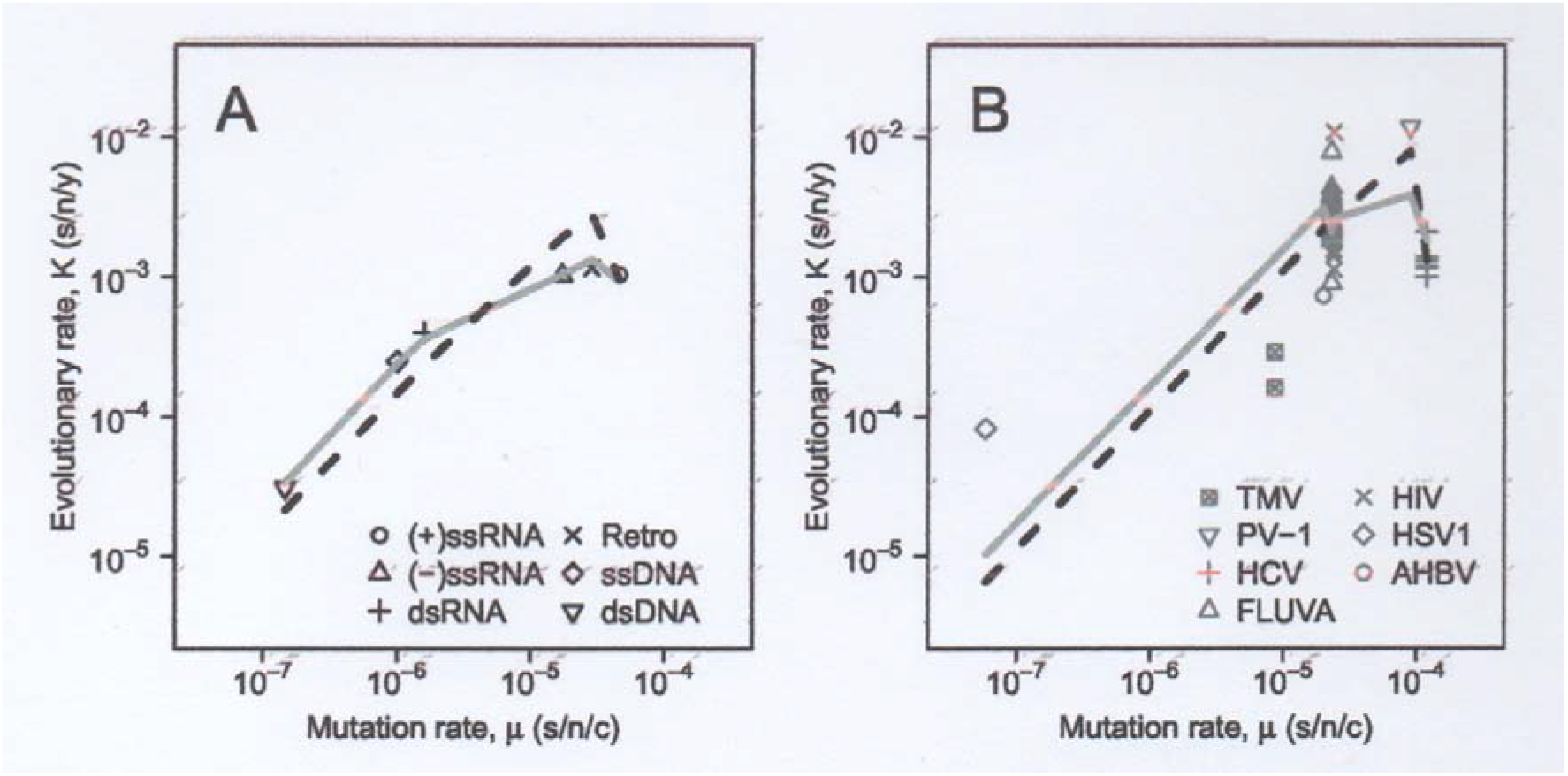
(A) Log-scale mean evolutionary rates against mutation rates for each Baltimore Class (B) Evolutionary rates against mutation rates for individual viruses. For both (A) and(B), the solid line represents the deleterious mutation model prediction, while the dashed line indicates the prediction from the within-host analytical model. After Peck et.al 2015.

**Fig 3:**
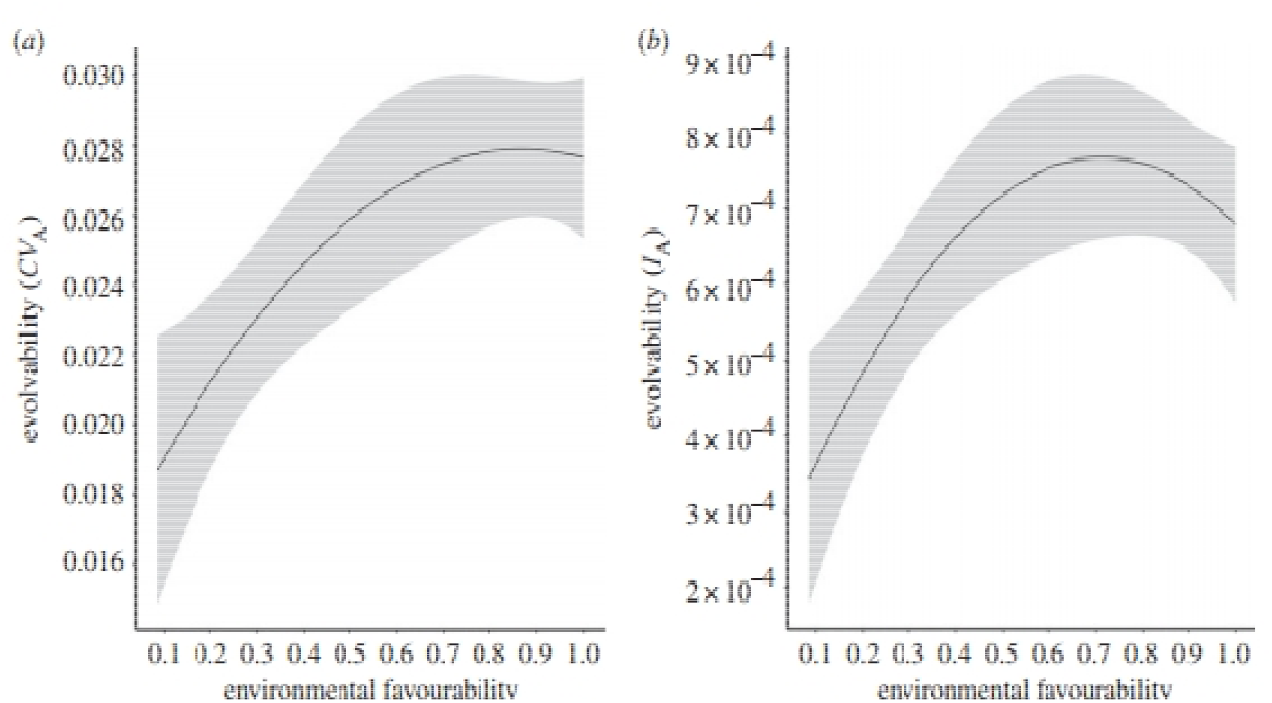
Evolvability criteria versus environmental favourability after Martenaz et.al 2017.

**Fig. 4.**
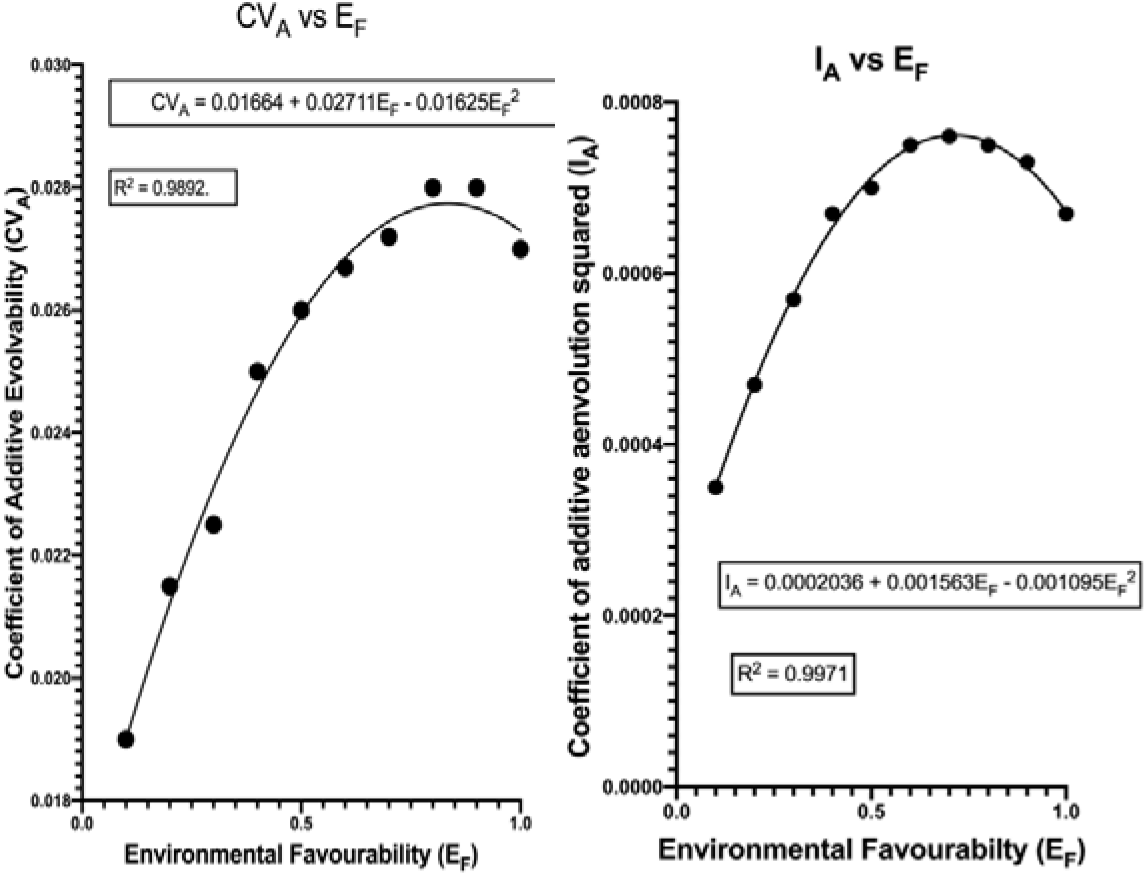
Coefficients against Environmental Favourability

Research into Covid-19 is obviously proceeding rapidly [Vilar & Isom 2020] and a good source of up-to-date information is the COG-UK website. Vilar & Isom’s analysis shows that SARS-CoV-2 proteins are mutating at substantially different rates, with most viral proteins exhibiting little mutational variability. The comprehensive temporal and geographical analyses showed two periods with different mutations in the SARS-CoV-2 proteome: December 2019 to June 2020 and July to November 2020. Some mutation rates differ also by geography; the main mutations in the second period occurred in Europe; this maybe a result of selection pressure. From the point of view of this theory a reference with a promising title is available [Pathana et.al.202]. However, this big data study records different mutation rates as percentage change, and the time base is the number of infections occurring over time, hence it is non-linear, and no environmental data was included; therefore, no direct comparisons can be made. However, the trends from their results show that mutations increase rapidly initially and then slow down over time. Recent research [Kemp et.al 2021] shows that a patient, with a compromised immune system being treated with plasma, demonstrated a considerable increase in mutation rates after the third infusion, which could correspond to this theory if the virus mutates rapidly when the environment becomes significantly adverse.

Dealing next with bacterial mutation rates, there is again a considerable literature [Rosche et.al 2000, Zeyl & Devisser 2000, Denamur & Matic 2006, Pal et.al. 2007 Wielgossa et.al 2013, Chevallereau et.al 2018, Ferenci 2019, Ramiro RS et.al 2020] however the same type of problem in comparing equation 8 with the evidence exists as does that of viral mutation rates; the mutation rates are not measured on a time base. One reference [Rosche et.al] discusses the problem with determining mutation rates in bacteria, and maintain that these methods are more accurate than time rate of change studies due to the need not to invoke theoretical models in the latter; however this should not preclude such studies. Rosche et.al 2000 surveyed six different methods of calculation of mutation rates and concluded that the maximum likelihood rate is preferred, but more work is necessary. Ferenci 2019 from a review concludes that recent findings suggest a very uneven relationship between environmental stress and mutations, and it remains to be investigated whether stress specific genetic variation impacts on evolvability differentially in distinct environments. Therefore, although again there do not appear to be any systematic investigations of bacterial mutation rates and environment, again there seems to be a general conclusion that bacterial mutation rates do respond to selective pressures from the environment, which could again be in agreement with the differential equation derived here.

Research into evolution at population level is extensive and has changed over time from qualitative to quantative measures, in some qualtative investigations it was difficult to separate phenotype plasticity from genetic evolvabilty. Following the landmark paper by Houle (1992) where he defined two quantative measures of genetic evolvablity namely the coefficient of evolvability CVA and its square I_A_, with the comment that he considered I_A_ to be the most accurate, the field became more quantative. Exploring genetic evolvability is complex and in the subsequent years there were many investigations which contained errors smmarised by Garcia-Gonzalez et.al. 2012 who reviewed 364 papers and found that there were errors in 47%.

In a major investigation of a 35 years longitudinal study Husby (2011) showed that, in a wild population of Great Tits (Parus major), the strength of the directional selection gradients on timing of breeding increased with increasing spring temperatures, and that genotype-by-environment interactions also predicted an increase in additive genetic variance, and heritability, of timing of breeding with increasing spring temperature. There was a significant positive association between the annual selection differentials and the corresponding heritability.

Marrot (2013) Posited that evolutionary adaptation, as a response to climate change, is expected for fitness related traits affected by climate and exhibiting genetic variance. Their results indicated an increase in the strength of selection by 46% for every +1°C anomaly in warmer daily maximum temperatures in April. Such climate-driven influences on the strength of directional selection acting on laying date could favour an adaptive response in this trait, since it is heritable.

Franks (2013) Observed that as climate change progresses, there are widespread changes in phenotypes in many plant populations. Whether these phenotypic changes are directly caused by climate change, and whether they result from phenotypic plasticity or evolution, are active areas of investigation. Of the 38 studies that met their criteria for inclusion, all found plastic or evolutionary responses, with 26 studies showing both.

Wadgyner et.al (2019) conducted a major review (192 references) of the effects of climate change on evolution and concluded that climate change has the potential to directly and indirectly influence the direction and rate of evolution in natural populations. They also considered coevolution with both species and the environment.

In another major investgation Martinez-Padilla et.al (2017) conducted a major bibliogaraphical review and found a final sample covering 20 populations of 12 species and asked the critical question: “do populations have the ability to evolve in response to these changes?”; however, knowledge on how evolution works in wild conditions under different environmental circumstances is extremely limited. They used published data to collect or calculate 135 estimates of evolvability of morphological traits of European wild bird populations. The authors used a quantative approach based on the two dimensionless ratios namely the coefficient of additive genetic variation (CV_A_), and its square (I_A_), as indexes of evolvability [Houle 1992]. If the time rate of change of evolution is proportianl to the environmental change in accordance with equation 8 the comparison with recent evidence of evolutionary change can be facilitated by rewriting equation 6 as follows: if a high rate of the coeficient of additive evolution (CV_A_) is due to an increase in the the rate of change of mutations, then it can be inferred that:

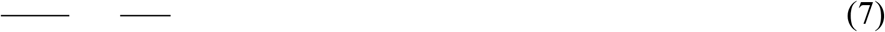

Where is the environmental favourability, and the dimensionless coefiicents defined after Houle 1992 as:

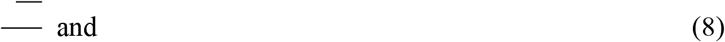

Where V_A_ ie the additive variation and the population mean.

The environmental favourability was determined using Logistic Regression methods, after Real et.al 2006, to obtain Favourability Functions.

Equation 7 makes it simpler to compare with the evidence presented below. The results see Fig 2 clearly show that the environmental favourability increases (fitness) the rate of evolvability (the slope of the curves) decreases; this is in accord with the differential equation of coevolution (equation 6).

Figure 2a shows that as the environmental favourabilty approaches 1.0 the rate of evolvabilty deceases to approximately zero, with a reasonably large but unambiguous range. In Figure 2b the results for I_A_ are more definite and show that as the envirommental favourability approaches 1.0 the rate of evolvabity actually becomes negative, which is understandable for if the fitness to environmental favourability is perfect why should the organism mutate? This is an important result as it as it shows that it is not evolution per.se which is driven by changes in the environment, but it is the rate of evolution (evlovability)which changes, which is in agrement with equation 6.

Curve fitting to the results above gives Fig 3a & 3b, from these figures it is obvious that the means are fitted very accurately by quadratic functions as shown.

Taking the analysis to the next step the rates of change for CA_V_ and I_A_ are:

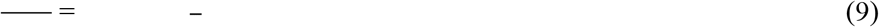

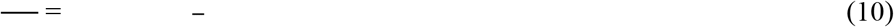

These equations, of the generic form A – BE_F_, clearly show that the slope of the CA_V_ and I_A_ consistently reduce as the environmental favourability increases, and equations 10 and 11 are linear functions of the environmental favourability. This could be important in helping to untangle the importance of the environmental changes and assist the definition of the Lagrangian in equation 6. In the face of widely held views on mass extinctions because of climate change this gives some small hope [Husby]. However, this is potentially bad news for Covid-19 as. although the data is not yet available, as the stress on the environment for the virus increases due to the vaccination programme, then the mutation rate will increase and we are in a continual catch-up, as with the influenza virus.

## Conclusions

Taking the environmental Coevolution model of Lewontin, a differential equation for Coevolution shows that it is the time rate of evolutionary change (evolvability) that responds to changes in the environment, which agrees with a number of studies. Further analysis shows that the rate of change of the coefficient of additive evolvability (Cav) and its square (IA) with environmental favourability are linear, which could help in defining the Lagrangian of the environmental effects. However, this is potentially bad news for Covid-19 as. although the data is not yet available, as the stress on the environment for the virus increases due to the vaccination programme, then the mutation rate will increase, and we are in a continual catchup scenario as with the influenza virus.

## Acknowledgement

The author acknowledges the contribution of Professor Ravindra Gupta, of the Cambridge Institute of Therapeutic Immunology and Infectious Disease.

## Notes

### Competing Interest Statement

The authors have declared no competing interest.

